# Occurrence of extended spectrum β-lactamase and AmpC-producing *Escherichia coli* in retail meat products from the Maritime Provinces, Canada

**DOI:** 10.1101/2020.08.25.266395

**Authors:** Babafela Awosile, Jessica Eisnor, Matthew E. Saab, Luke Heider, J T. McClure

**Affiliations:** Health Management, University of Prince Edward Island, 550 University Avenue, Charlottetown, C1A 4P3, Prince Edward Island, Canada

**Keywords:** Antimicrobial resistance, Extended-spectrum cephalosporins, β-lactamases, *Escherichia coli*, Retail meat products

## Abstract

This study was conducted to determine the occurrence of antimicrobial resistance to the extended-spectrum cephalosporins (ESC) in *Escherichia coli* isolates recovered from retail meat products collected in the Maritime Provinces of Canada using both selective and traditional culture methods, and genotypically using multiplex polymerase chain reactions.

ESC-R *E. coli* was detected in 33/559 (5.9%) samples using the traditional culture compared to 151/557 (27.1%) samples using the selective culture method. The recovery of ESC-R *E. coli* isolates was more common in poultry compared to beef and pork (P<0.001). Multi-drug resistance, ESBL, and AmpC phenotypes were more common in chicken-derived isolates than other retail meat products (P<0.001). From the 98 isolates selected, 76 (77.6%) isolates were positive for either ESBL and AmpC β-lactamases or both. Among the 76 isolates, *bla*_CMY-2_ (78.9%), *bla*_CTXM_ (46.1%), *bla*_TEM_ (21.1%), and *bla*_SHV_ (1.3%) were detected. Among the *bla*_CTXM_-producing isolates; *bla*_CTXM-1_, *bla*_CTXM-2_, and *bla*_CTXM-9_ phylogenetic groups were detected. β-lactamase genes were detected more in chicken-derived isolates compared to other meat types (P<0.01). This study demonstrated the occurrence of ESBL and AmpC resistance genes in retail meat products in Maritime Provinces of Canada. Also, selective culture significantly improved the recovery of ESC-R *E. coli* isolates from retail meat samples.

## Introduction

Zoonotic foodborne pathogens, including antimicrobialresistant (AMR) bacteria, can be transmitted from animals to humans through the food chain. AMR bacteria may be transferred to retail meat products during slaughtering, processing, and subsequent handling of meat products. Fecal contamination at slaughter is the primary source of contamination although personnel or environmental contamination may occur. Contamination of retail meat products is a potential source of AMR bacterial exposure to humans through consumption (Nekouei et al. 2018). AMR bacteria of importance include extended-spectrum cephalosporin (ESC)resistant *Salmonella* spp. and *Escherichia coli*, as well as fluoroquinolone-resistant *Campylobacter* spp. in retail beef, chicken, turkey, and pork products and the processing plants (EFSA Panel on Biological Hazards (BIOHAZ) 2011). Infections caused by AMR pathogens areassociated with prolonged duration of illness, bloodstream infections, increased health care costs, prolonged hospitalization, and increased mortality(Angulo et al. 2004).

World Health Organization and Health Canada’s Veterinary Drug Diretorate classify extended-spectrum (third and fourth generation) cephalosporins as highly important antimicrobials for human medicine (World Health Organization 2014; Ebrahim et al. 2016). Increased resistance to this category of drugs has been reported, mainly mediated by β-lactamase genes, such as extended-spectrum β-lactamases (ESBL) and AmpC β-lactamases (Carattoli 2008; Smet et al. 2010). This increased resistance is partly due to the ease of acquisition and dissemination of antimicrobial resistance genes in commensal bacteria, especially in *E. coli* (Seiffert et al. 2013). *Escherichia coli* is prevalent in the gastrointestinal tracts of animals and humans; therefore, they are commonly used as an indicator organism for fecal contamination and sentinel for the surveillance and monitoring of antimicrobial resistance (EFSA Panel on Biological Hazards (BIOHAZ) 2011). Different studies have revealed that the use of ceftiofur, the only ESC approved for use in food animals in North America, is associated with the recovery of both AmpC and ESBL-producing *E. coli* isolates in food animals (Daniels et al. 2009; Schmidt et al. 2013; Saraiva et al. 2018). Similarly, β-lactamase producing *E. coli* isolates have also been reported in retail meat products (Zhao et al. 2012; Sheikh et al. 2012) and are a reservoir for ESBL and AmpC producing bacteria for human exposure. A study within Canada has reported a strong correlation between ceftiofur-resistant *Salmonella enterica* serovar Heidelberg isolated from retail chicken and incidence of ceftiofur-resistant *S*. Heidelberg infections in humans across Canada (Dutil et al. 2010). There is a need for continuous surveillance of ESC-resistance and associated β-lactamase resistance genes to better understand the risk of exposure to humans from to contaminated retail meat products.

In Canada, AMR monitoring and surveillance in retail meat products is coordinated by the Canadian Integrated Program for Antimicrobial Resistance Surveillance (CIPARS) (Government of Canada, 2015). This routine surveillance is used to generate information for measuring the risk of human exposure to AMR bacteria associated with the consumption of retail meat products. This information is based on non-selective culture of bacterial organisms and antimicrobial susceptibility testing, with limited reports on the molecular basis of resistance. In addition, it has been hypothesized that use of non-selective, traditional culture methodology underestimates the recoveries and frequencies of resistant-bacteria in the laboratory (Dutil et al. 2010). Apart from routine monitoring by CIPARS, a previous molecular study on ESC-resistant *E. coli* within Maritime Provinces, Canada was based on the recovery of ESC-resistant *E. coli* (ESC-R *E*. coli) and *Salmonella* spp. in marketed retail meat products from Nova Scotia using conventional culture method (Forward et al. 2004). From that study, the *bla*_CMY-2_ gene was detected in the retail meat products.

In this study, our first objective was to compare the frequency of recovery of ESC-R *E. coli* from retail meat products using both selective and traditional culture methodologies. We hypothesize that using ESC selective culture methods would enhance recovery of ESC-R *E. coli* from retail meat samples and that we would detect both ESBL and AmpC resistant genes from these isolates. Our second objective was to estimate the prevalence and determine the antimicrobial susceptibility patterns of ESC-R *E. coli* recovered using selective culture from retail meat products collected in the Maritime Provinces, Canada. Our last objective was to examine the molecular basis of ESC resistance in a selected ESC-R *E. coli* isolates recovered from the retail meat products.

## Materials and methods

### Retail meat samples collection

Bone-in, skin-on chicken pieces, ground beef, pork chops, and turkey (ground or bone-in, skin-on pieces) were purchased from grocery stores, independent markets, or butcher shops. Retail meat samples of different package sizes, i.e., regular or family size were sampled. Samples were randomly collected in the Maritime Provinces (New Brunswick, Nova Scotia, and Prince Edward Island) in a stratified-multistage hybrid design where selection of census divisions within a province for sample collections is weighted relative to the population. Sample collection was carried out from June to December 2013 as part of the retail surveillance component of CIPARS. Two census divisions (counties) were sampled each week, and four stores were sampled in each division, with at least one store being an independent market or butcher shop. One sample each from pork, chicken, turkey, and beef meat products was purchased from each store. Once purchased, samples were placed on ice in coolers and returned to the laboratory at the Atlantic Veterinary College where they were held at 5°C ± 1 for processing the following day.

### Isolation and identification of ESC-R *E. coli*

From the meat samples, one piece of bone-in, skin-on chicken or turkey, one pork chop, 25±1 g of ground beef or ground turkey were used for the isolation and identification of ESC-R *E. coli*. Initial pre-enrichment was carried out by rinsing the meat samples in buffered peptone water (BPW) and placing the meat samples in the BPW on an orbital shaker for 10 minutes. The BPW-meat rinsate was placed in equal volume (1:1) into double strength of *E. coli* broth (EC broth) and incubated overnight at 44°C for 18-24 hours. For traditional culture method, 10μl. of sample-EC broth was plated onto an eosin methylene blue (EMB) agar plate. Plates were then incubated for 18-24 hours at 35°C± 1°C.

A tryptic soy agar plate with 5% sheep blood (BTSA) containing vancomycin (6 μg/mL), amphotericin B (2 μg/mL), ceftazidime (2 μg/mL), and clindamycin (1 μg/mL) (VACC) was used for selective culture of ESC-R *E. coli* (Singh et al. 2012). VACC plates were inoculated with 50 μL of sample-EC broth and incubated at 35°C for 18-24 hours. VACC plates with no presumptive *E. coli* colonies after 24 hours were incubated for an additional 18-24 hours. Presumptive *E. coli* colonies were sub-cultured to eosin methylene blue agar, and typical colonies were purified on a BTSA. *E. coli* isolates were confirmed using biochemical tests including lactose-fermentation, utilization of citrate, and indole test. All the isolates were frozen in Brucella broth with 15% glycerol at −80°C for further laboratory analysis.

### Antimicrobial susceptibility testing

Antimicrobial susceptibility testing was performed all the isolates using the Sensititre™ broth microdilution systemto determine minimum inhibitory concentrations (MICs). Testing was performed according to the Clinical and Laboratory Standards Institute guidelines (CLSI, 2013). The CMV2AGNF (Sensititre™, Trek™ Diagnostic Systems, Westlake, Ohio) susceptibility plate of the National Antimicrobial Monitoring System (NARMS) containing 14 antimicrobials was used in this study. However, azithromycin was excluded from analysis because of the intrinsic resistance in *E. coli*. The following antimicrobial agents were tested with the resistance breakpoints presented in parentheses (CLSI, 2013): ampicillin (≥32 μg/ml), amoxicillin-clavulanic acid (AMC, ≥32/16 μg/ml), chloramphenicol (≥32 μg/ml), ceftriaxone (≥4 μg/ml), ceftiofur (≥8 μg/ml), ciprofloxacin (≥4 μg/ml), cefoxitin (≥32 μg/ml), gentamicin (≥16 μg/ml), kanamycin (≥64 μg/ml), nalidixic acid (≥32 μg/ml), streptomycin (≥64 μg/ml), sulfisoxazole (≥512 μg/ml), trimethoprim-sulfamethoxazole (TMS, ≥4/76 μg/ml), and tetracycline (≥16 μg/ml). Multi-drug resistance was based on the World Health Organization’s definition of resistance to at least one antimicrobial each in ≥3 antimicrobial classes (Magiorakos et al. 2012). Based on MIC testing, isolates showing resistance to ceftriaxone and/or ceftiofur were considered as ESBL phenotypes. While isolates demonstrating resistance to cefoxitin, amoxicillin-clavulanate, in addition to resistance to ceftriaxone and/or ceftiofur, were considered as an AmpC phenotypes.

### Molecular detection of β-lactamase resistance genes

Genomic DNA of presumptive ESC-R *E. coli* isolates was extracted using the InstaGene™ Matrix following manufacturer’s guidelines (Bio-Rad, Montreal Canada). For all multiplex PCR assays, the Qiagen multiplex PCR kit (Qiagen, Mississauga, Ontario, Canada) with 1X Qiagen multiplex PCR master mixture, 1X Q-solution, and molecular grade water, together with 1X consensus primer pair mixture as well as 2 μl of template DNA were included in the final 25 μl mixture, according to the manufacturer’s instructions. Positive and negative controls were included in every multiplex PCR. Amplification and DNA fragment electrophoresis was carried out as previously described and reported (Awosile et al., 2018).

### Statistical analysis

Comparison of frequency of recovery of ESC-R *E. coli*, as well as the frequency of antimicrobial resistance between the meat products, was carried out using the Chi-squared test or Fisher’s exact test. McNemar’s chi-squared test was used to compare the proportion of ESC-R *E. coli* between culture methods. Further analysis was done on the ESC-R *E. coli* recovered through selective culture medium. Association between various data collected and recovery of ESC-R *E. coli* was explored using multivariable logistic regression model. Initial univariable unconditional association (P<0.25) was explored between the recovery of ESC-R *E. coli* and type of retail meat products (chicken, beef, pork, or turkey), provinces where meat samples were collected, store operation types (butcher, independent or chain), whether meat products were packaged in store or not, and retail meat size (regular size or family pack). Due to the hierarchical nature of the data, a mixed effect logistic model was carried out to account for clustering. However, the random effects of stores, census division and province were non-significant. Therefore, multivariable logistic regression was carried out to explore the relationship between the independent variables and the outcome of interest. We used stepwise model selection procedure to further select the predictors into the final multivariable model despite the initial univariable model. The interaction between the predictors was explored. Likelihood ratio test was used to examine the model adequacy and comparison. We selected the best model based on low Akaike’s information criterion. The Wald test was used to test for the significance of each predictor in the model and level of statistical significance of predictors was considered at P<0.05. All statistical analysis was done using Stata 15 (IC Stata Corp, College Station, Texas, USA).

## Results

A total of 559 raw retail meat samples (144 chicken, 144 beef, 144 pork and 127 turkey) were collected from June to December 2013. ESC-R *E. coli* was detected in 33/559 (5.9%, 95% CI: 3.9%-7.9%) samples using the traditional culture method and 151/557 (27.1%, 95% CI: 23.4%-30.8%) samples using the selective culture method (Table 1, Figure 1). Two isolates cultured using the selective method did not undergo MIC testing. Selective culture detected a significantly higher proportion of ESCR *E. coli* compared to the traditional method (*p*<0.001). The selective method was able to detect more ESCR *E. coli* than the traditional method for all four commodities tested. There was a significant difference in the ESCR *E. coli* frequency between the meat commodities using both traditional and selective culture methods (*p*<0.001 for both methods). When comparing the prevalence of ESC-R *E. coli* isolated between commodities, chicken samples had the highest apparent prevalence of ESC-R *E. coli* using traditional culture methods (18.8%, 27/144) when compared to beef, pork and turkey commodities (Figure 1). The recovery of ESC-R *E. coli* increased in all commodities using the selective culture methods, especially chicken products (65.3%, 94/144) (Figure 1).

**Table 1.** Comparison of ESC-R *E. coli* recovery between selective and traditional methodologies.

|  |  | Selective Method |  |  |
| --- | --- | --- | --- | --- |
|  |  | Detected | Not detected | Total |
| **Traditional Method** | Detected | 31 | 2 | 33 |
|  | Not detected | 120 | 404 | 524 |
|  | Total | 151 | 406 | 557 |

McNemar's test ( $p < 0.001$ )

**Figure 1.**
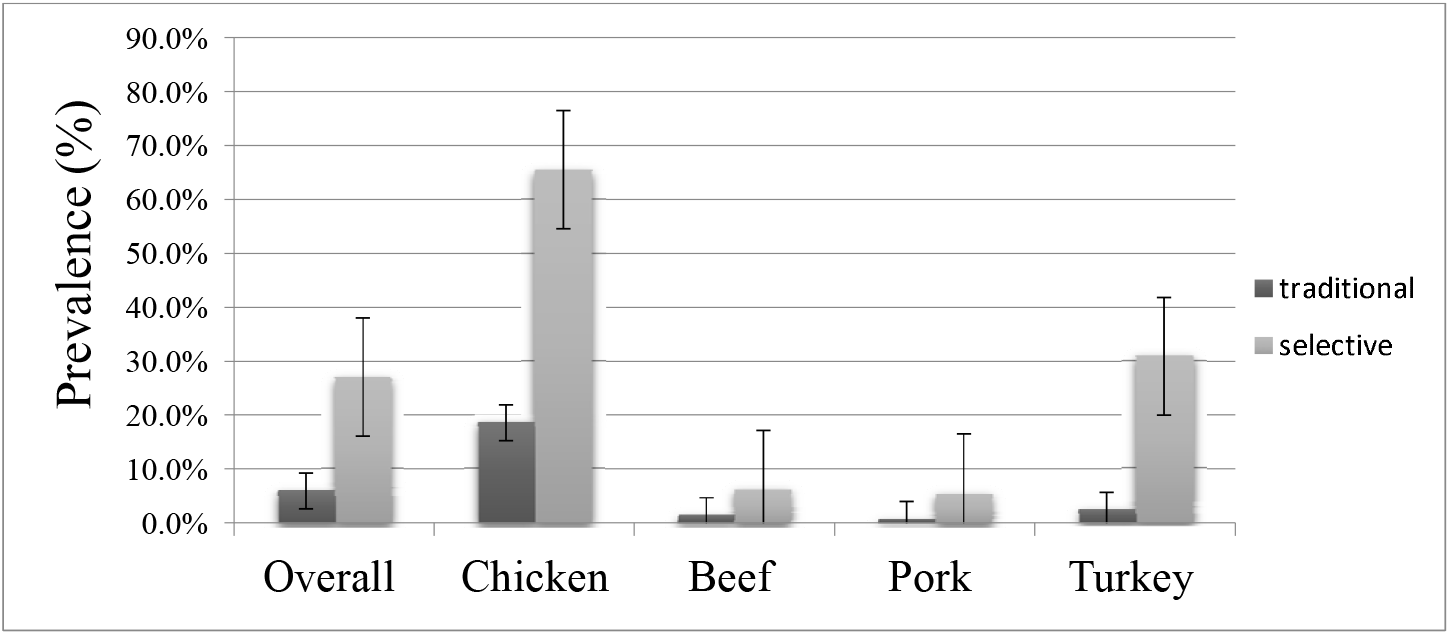
ESC-R *E. coli* prevalence estimates with standard errors in retail meat samples using both selective and traditional culture methodologies.

For the isolates recovered from the selective culture medium, the frequency of recovery of ESC-R *E. coli* isolates based on retail meat types, store operation type, provinces, retail meat package size, and in-store processing were presented in Table 2. The recovery of ESC-R *E. coli* isolates was more common in poultry compared to beef and pork. More ESC-R *E. coli* isolates were recovered from the in store processed meat products than from meat products processed elsewhere. There was a trend between recovery of ESC-R *E. coli* isolates and the retail meat size package, with large “family packs” having more recovery of ESC-R *E. coli*. While frequency of ESC-R *E. coli* recovery between the three provinces (26-28%) and between the store operation types ware almost similar (26-28%).

**Table 2.** Prevalence and final multivariable logistic regression of factors associated with ESC-R *E. coli* recovered using selective culture from retail meat products from the Maritime Provinces, Canada

| Variables | Categories | Number of samples | Prevalence |  | Multivariable model |  |  |  |
| --- | --- | --- | --- | --- | --- | --- | --- | --- |
|  |  |  | % | 95% CI | Coeff | Odds ratio | 95% CI | P-value |
| Retail meat types | Beef | 143 | 6.29 | 3.29-11.69 | Reference | - | - | - |
|  | Chicken | 144 | 65.28 | 57.11-72.64 | 3.15 | 23.57 | 10.88-51.04 | <0.001 |
|  | Pork | 143 | 5.59 | 2.81-10.83 | -0.09 | 0.91 | 0.34-2.43 | 0.847 |
|  | Turkey | 127 | 31.50 | 23.98-40.12 | 1.45 | 4.25 | 1.69-10.71 | 0.002 |
| Store operation type | Butcher | 71 | 27.14 | 17.96-38.80 |  |  |  |  |
|  | Chain | 430 | 26.80 | 22.81-31.21 |  |  |  |  |
|  | Independent | 56 | 28.57 | 18.20-41.82 |  |  |  |  |
| Provinces | New Brunswick | 279 | 28.05 | 23.07-33.65 |  |  |  |  |
|  | Nova Scotia | 248 | 26.20 | 21.09-32.06 |  |  |  |  |
|  | Prince Edward Island | 30 | 24.14 | 11.79-43.10 |  |  |  |  |
| In store processing | No | 171 | 43.86 | 36.57-51.41 | Reference | - | - | - |
|  | Yes | 385 | 19.58 | 15.89-23.87 | -0.59 | 0.55 | 0.29-1.02 | 0.059 |
| Retail meat size | Family pack | 32 | 46.87 | 30.32-64.15 |  |  |  |  |
|  | Regular | 525 | 25.81 | 22.23-29.75 |  |  |  |  |

From the initial univariable logistic regression model, package size and within store processing were unconditionally associated (P<0.25) with the recovery of ESC-R *E. coli* isolates from retail meats products. From the stepwise model selection procedure, retail meat type, package size, store operation type and within store processing were selected for the initial multivariable model. However, retail meat type (P<0.001) was statistically associated with the recovery of ESC-R- *E. coli* in the final multivariable regression model (Table 2). The recovery of ESC-R *E. coli* isolates was 23.56 times more likely from chicken meat products than from beef products (reference) (95% CI: 10.88-51.04, P<0.001). The recovery of ESC-R *E. coli* isolates was 4.25 times more likely from turkey meat products than from beef products (95% CI: 1.69-10.71, P=0.002). There was no significant difference in the recovery of ESC-R- *E. coli* isolates when comparing pork samples to the recovery in beef samples.

A higher proportion of AMR (89-100%) was seen among the ESC-R *E. coli* isolates to ampicillin, amoxicillin-clavulanate, ESCs, and cefoxitin (Table 3). A low level of resistance to kanamycin (15.89%) and gentamicin (31.79%) was observed, and over half of the isolates were resistant to streptomycin (46.25%), sulfisoxazole (56.95%), and tetracycline (56.95%). The isolates were commonly susceptible to ciprofloxacin, nalidixic acid, chloramphenicol, and TMS. Ninety-four percent (94%) of ESC-R *E. coli* isolates showed β-lactam resistance consistent with AmpC phenotypes (Table 3). Chicken-derived ESC-R *E. coli* isolates showed more resistance to almost all the antimicrobials tested compared to other retail meat types (Table 3). Multi-drug resistance (MDR) was seen in 94.7% of the ESC-R *E. coli* isolates, while more MDR ESC-R *E. coli* isolates were also recovered from chicken meat products compared to the other retail meat types. Both ESBL and AmpC phenotypes were isolated more from chicken meat products compared to other meat products tested. Among the isolates, 44 MDR patterns were observed (Table 4). Co-resistance to streptomycin, tetracycline, and sulfisoxazole was commonly observed with either ESBL or AmpC phenotypes or with both.

**Table 3.** Antimicrobial resistance profile in all the ESC-R *E. coli* isolates (n=151) recovered from retail meat products from the Maritime Provinces, Canada

| Antimicrobials | Number of resistant isolates<br>(n) | Retail meat products |  |  |  | P-value | Total resistant (%) |
| --- | --- | --- | --- | --- | --- | --- | --- |
|  |  | Beef (%) | Chicken (%) | Pork (%) | Turkey (%) |  |  |
| *Ampicillin | 151 | 5.96 | 62.91 | 5.30 | 25.83 | <0.001 | 100 |
| *AMC | 148 | 6.08 | 64.19 | 5.41 | 24.32 | 0.032 | 98.01 |
| *Chloramphenicol | 15 | 20.00 | 80.00 | 0.00 | 0.00 | 0.007 | 9.93 |
| *Ceftriaxone | 151 | 5.96 | 62.91 | 5.30 | 25.83 | <0.001 | 100 |
| *Ceftiofur | 135 | 5.19 | 65.93 | 2.96 | 25.93 | 0.001 | 89.40 |
| *Cefoxitin | 142 | 4.23 | 66.90 | 2.82 | 26.06 | <0.001 | 94.03 |
| Ciprofloxacin | 5 | 0.00 | 60.00 | 0.00 | 40.00 | 0.799 | 3.31 |
| Gentamicin | 48 | 2.08 | 72.92 | 0.00 | 25.00 | 0.082 | 31.79 |
| Kanamycin | 24 | 8.33 | 45.83 | 4.17 | 41.67 | 0.171 | 15.89 |
| Nalidixic acid | 5 | 0.00 | 60.00 | 0.00 | 40.00 | 0.799 | 3.31 |
| Streptomycin | 70 | 7.14 | 64.29 | 1.43 | 27.14 | 0.249 | 46.35 |
| Sulfisoxazole | 86 | 5.81 | 61.63 | 3.49 | 39.07 | 0.550 | 56.95 |
| *Tetracycline | 86 | 5.81 | 55.81 | 2.33 | 36.05 | 0.005 | 56.95 |
| TMS | 19 | 15.79 | 63.16 | 5.26 | 15.79 | 0.201 | 12.58 |
| *ESBL phenotype | 151 | 5.96 | 62.91 | 5.30 | 25.83 | <0.001 | 100 |
| *AmpC phenotype | 142 | 4.22 | 62.91 | 2.65 | 24.50 | <0.001 | 94.04 |
| *MDR isolates | 143 | 4.90 | 65.73 | 2.80 | 26.57 | <0.001 | 94.7 |

\*Statistically significant recovery between meat commodities at P<0.05
AMC-Amoxicillin-clavulanate, TMS-Trimethoprim-sulfamethoxazole, MDR- Multidrug resistance

**Table 4.** Multi-drug resistance patterns in ESC-R- *E. coli* (n=151) recovered from retail meat products from the Maritime Provinces, Canada

| Resistance patterns | Frequency | % |
| --- | --- | --- |
| FoxCefCaxAgAmp | 33 | 21.85 |
| StrTetSulGmFoxCefCaxAgAmp | 16 | 10.6 |
| StrTetSulKanFoxCefCaxAgAmp | 9 | 5.96 |
| StrTetSulTmsFoxCefCaxAgAmp | 9 | 5.96 |
| TetFoxCefCaxAgAmp | 9 | 5.96 |
| StrTetSulGmFoxCefCaxCAgAmp | 7 | 4.64 |
| SulFoxCefCaxAgAmp | 7 | 4.64 |
| StrSulGmFoxCefCaxAgAmp | 6 | 3.97 |
| CaxAgAmp | 5 | 3.31 |
| TetSulGmFoxCefCaxAgAmp | 5 | 3.31 |
| FoxCaxAgAmp | 3 | 1.99 |
| StrTetKanFoxCefCaxAgAmp | 3 | 1.99 |
| StrTetSulGmKanFoxCefCaxCAgAmp | 3 | 1.99 |
| GmFoxCefCaxAgAmp | 2 | 1.32 |
| NalFoxCipCefCaxAgAmp | 2 | 1.32 |
| StrTetFoxCefCaxAgAmp | 2 | 1.32 |
| TetSulFoxCefCaxAgAmp | 2 | 1.32 |
| SulGmFoxCefCaxAgAmp | 2 | 1.32 |
| CefCaxAgAmp | 1 | 0.66 |
| StrFoxCefCaxAgAmp | 1 | 0.66 |
| StrKanFoxCefCaxAgAmp | 1 | 0.66 |
| StrTetSulFoxCaxAgAmp | 1 | 0.66 |
| StrTetSulFoxCaxCAgAmp | 1 | 0.66 |
| StrTetSulFoxCefCaxAgAmp | 1 | 0.66 |
| StrTetSulGmFoxCaxAgAmp | 1 | 0.66 |
| StrTetSulGmKanCaxAmp | 1 | 0.66 |
| StrTetSulGmKanFoxCefCaxAgAmp | 1 | 0.66 |
| StrTetSulNalGmFoxCipCefCaxCAgAmp | 1 | 0.66 |
| StrTetSulTmsFoxCefCaxCAgAmp | 1 | 0.66 |
| StrTetSulTmsGmFoxCefCaxAgAmp | 1 | 0.66 |
| StrTetSulTmsGmKanFoxCefCaxCAgAmp | 1 | 0.66 |
| StrTetSulTmsKanFoxCefCaxAgAmp | 1 | 0.66 |
| StrTetSulTmsKanFoxCefCaxCAgAmp | 1 | 0.66 |
| StrTetSulTmsNalFoxCipCefCaxAgAmp | 1 | 0.66 |
| TetKanFoxCefCaxAgAmp | 1 | 0.66 |
| TetNalKanFoxCipCaxAmp | 1 | 0.66 |
| TetTmsFoxCefCaxAgAmp | 1 | 0.66 |
| TetSulCefCaxAmp | 1 | 0.66 |
| TetSulFoxCaxAgAmp | 1 | 0.66 |
| TetSulGmFoxCaxAgAmp | 1 | 0.66 |
| TetSulKanCefCaxAgAmp | 1 | 0.66 |
| TetSulTmsFoxCefCaxAgAmp | 1 | 0.66 |
| SulTmsFoxCaxAgAmp | 1 | 0.66 |
| SulTmsFoxCefCaxAgAmp | 1 | 0.66 |
| Total | 151 | 100 |

Ag- Amoxicillin-clavulanate, Amp- Ampicillin, Cef- Ceftiofur, Cax- Ceftriaxone, C- Chloramphenicol, Str- Streptomycin, Sul- Sulfisoxazole, Tms- Trimethoprim-sulphamethoxazole, Nal- Nalidixic acid, Tet- Tetracycline

Four different types of β-lactamase genes were detected among the isolates: *bla*_CMY-2_, *bla*_TEM_, *bla*_SHV_, and *bla*_CTXM_ (Table 5). From the 98 isolates selected, 76 (77.6%) were positive for either ESBL and/or AmpC β-lactamases, while 22 isolates were negative for the β-lactamase resistance genes tested. Among the 76 isolates positive for at least one β-lactamase gene, *bla*_CMY-2_ (78.9%) was most detected, followed by *bla*_CTXM_ (46.1%), *bla*_TEM_ (21.1%) and then one isolate was positive for *bla*_SHV_ (1.3%). Among the *bla*_CTXM_-producing isolates (n=35), three different phylogenetic groups were detected, and these included *bla*_CTXM-1_(42.9%), *bla*_CTXM-2_, (40%), and *bla*_CTXM-9_ (17.1%). β-lactamase genes (*bla*_CMY-2_, *bla*_TEM_ and *bla*_CTXM_) were detected more in poultry-derived isolates (chicken or turkey) compared to other meat types (P<0.01). Presence of two or more β-lactamase resistance genes was detected in 35.7% of the isolates (Table 6), with 13.2% of the isolates carrying both *bla*_CMY-2_ and *bla*_CTXM-2_ genes. Among the 22 isolates that were negative for the tested β-lactamase genes, 11 isolates were phenotypically ESBL. Nine of these 11 isolates also showed AmpC phenotypic characteristics, while the other 11 isolates showed β-lactamase inhibitor resistance.

**Table 5.**
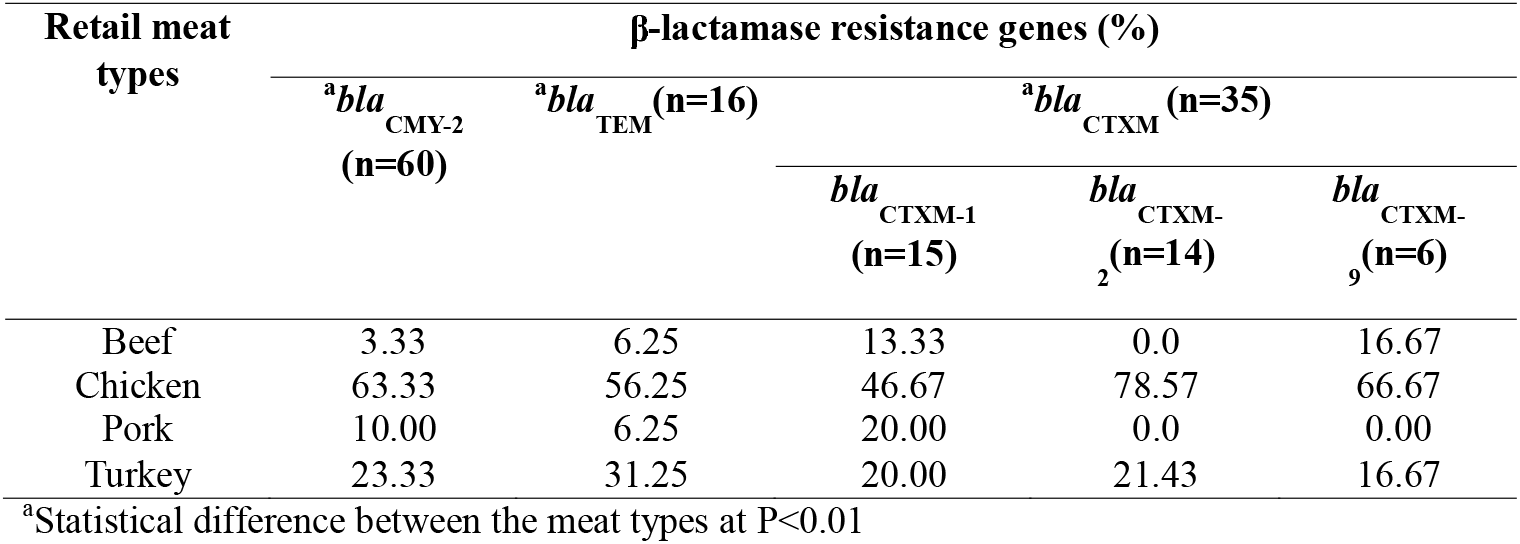
β-lactamase resistance genes in the selected ESC-R *E. coli* isolates (n=76) recovered from retail meat products from the Maritime Provinces, Canada

**Table 6.** β-lactamase genes patterns in selected ESC-R *E. coli* isolates (n=76) recovered from the retail meats from the Maritime Provinces, Canada

|  | **Patterns** | **β-lactamase genotypes** | **Frequency (%)** |  |
| --- | --- | --- | --- | --- |
| 1 | *bla*<sub>CMY-2</sub> | AmpC | 30 | 39.5 |
| 2 | *bla*<sub>CMY-2</sub> *bla*<sub>CTXM-2</sub> | ESBL, AmpC | 10 | 13.2 |
| 3 | *bla*<sub>CMY-2</sub> *bla*<sub>CTXM-1</sub> | ESBL, AmpC | 9 | 11.8 |
| 4 | *bla*<sub>TEM</sub> | ESBL<sup>a</sup> | 7 | 9.2 |
| 5 | *bla*<sub>CMY-2</sub> *bla*<sub>CTXM-9</sub> | ESBL, AmpC | 5 | 6.6 |
| 6 | *bla*<sub>CMY-2</sub>*bla*<sub>TEM</sub> | ESBL, AmpC | 4 | 5.3 |
| 7 | *bla*<sub>CTXM1</sub> | ESBL | 4 | 5.3 |
| 8 | *bla*<sub>CMY-2</sub> *bla*<sub>CTXM-1</sub>*bla*<sub>TEM</sub> | ESBL, AmpC | 1 | 1.3 |
| 9 | *bla*<sub>CMY-2</sub> *bla*<sub>CTXM-2</sub>*bla*<sub>TEM</sub> | ESBL, AmpC | 1 | 1.3 |
| 10 | *bla*<sub>CTXM-2</sub> | ESBL | 1 | 1.3 |
| 11 | *bla*<sub>SHV</sub> *bla*<sub>CTXM-2</sub> | ESBL | 1 | 1.3 |
| 12 | *bla*<sub>CTXM-1</sub>*bla*<sub>TEM</sub> | ESBL | 1 | 1.3 |
| 13 | *bla*<sub>CTXM-2</sub>*bla*<sub>TEM</sub> | ESBL | 1 | 1.3 |
| 14 | *bla*<sub>CTXM-9</sub>*bla*<sub>TEM</sub> | ESBL | 1 | 1.3 |

<sup>a</sup>Could be narrow-spectrum β-lactamase, ESBL, or inhibitor resistant *bla*<sub>TEM</sub>

## Discussion

The current study determined that the use of a selective medium does significantly increase the recovery of ESC-R *E. coli* from retail meat samples. From this study, the prevalence estimates of ESC-R *E. coli* increased significantly using the selective media. This finding provides evidence that traditional culture methodology may not be a reliable means of isolating and identifying ESC-R *E. coli* from retail meat samples. It also suggests that use of traditional culture methods for large scale surveillance such as CIPARS may not be accurately identifying the prevalence of AMR bacteria in retail meat samples. Previous studies have also arrived at a similar conclusion including studies on the recoveries of Shiga-toxin producing *E. coli*, ESC-R *E. coli* and *Salmonella* spp. (Dutil et al. 2010; Gill et al. 2014). It is possible for a sample to contain both ESC susceptible and ESC resistant isolates of the same species(Rawat and Nair 2010). In that case, the ESC resistant cells will likely be present in low numbers and may, therefore, frequently be missed especially when there are no morphological differences between ESC-R and susceptible colonies. When using a selective medium, this is not an issue because it is assumed that any growth on the plate will be resistant to the antimicrobial contained within the medium. Selective medium is superior at ESC-R *E. coli* detection when compared to traditional culture methods.

To understand the epidemiology of ESC R *E. coli* from retail meat products, it is important to know different β-lactamase resistance genes associated with the occurrence of extended-spectrum resistance. This study was also conducted to provide this information in *E. coli* isolated from retail meat products within the Maritime Provinces of Canada. Previous study on ESC-resistance in retail meat products within this region provided data for just one province (Nova Scotia) and also only reported on the detection of the the *bla*_CMY-2_ gene in the samples collected(Forward et al. 2004). Furthermore, surveillance reports by CIPARS only provide information on the prevalence and antimicrobial susceptibility data for ESC-resistant *E. coli* isolates from retail meat products with no data on associated resistance genes. In the present study, we demonstrated the occurrence of ESBL and AmpC resistance genes in retail meat products in all three Maritime Provinces of Canada. We observed a similar pattern of *E. coli* recovery from retail meat commodities in our study to studies from Western Canada and USA, as well as the previous study within Maritimes, Canada (Forward et al. 2004; Sheikh et al. 2012; Zhao et al. 2012; Yu et al. 2015) as *E. coli* recovery was greatest from chicken products, followed by turkey, beef, and lastly pork. This finding was consistent with the multivariable model of this study in which the odds of recovery of ESC-R *E. coli* was more likely in poultry meat products compared to other meat products. This finding further supports the report that placed poultry as one of the major source of foodborne bacteria that causes most of the foodborne outbreaks in North America (Chai et al. 2017).

Most of the ESC-R *E. coli* isolates were MDR including AMR to streptomycin, sulfisoxazole and tetracycline commonly observed, but these isolates had a low prevalence of AMR to aminoglycosides, quinolones, chloramphenicol, and TMS. This is a similar pattern to what has been reported by previous studies (Cook et al. 2009; Sheikh et al. 2012; Zhao et al. 2012; Yu et al. 2015). Co-resistance to streptomycin, sulfisoxazole, and tetracycline is commonly associated with ESBL and AmpC phenotypes, especially in MDR bacteria that harbor mobile genetic elements, including plasmids and integrons, that are capable of acquiring and disseminating AMR genes, such as *E. coli* (Seiffert et al. 2013). Variation in AMR profile was observed in the ESC-R *E. coli* isolates depending on the retail meat types. Among the 14 antimicrobials tested in this study, chicken-derived *E. coli* isolates demonstrated higher frequency of AMR to all the antimicrobials (including the clinically important antimicrobials) than turkey, beef, and pork. Similar patterns were also observed for the ESBL, AmpC and MDR phenotypic characteristics, as the proportions were higher in poultry-derived isolates than other meat types. Similar AMR patterns between the retail meat products have been reported from Western Canada and USA (Zhao et al. 2012; Sheikh et al. 2012).The variation in AMR patterns between retail meat products may be a reflection of the selective pressure created by antimicrobial usage pattern and management practices in different food animal productions and processing. Antimicrobial use either for prophylactic and/or therapeutic purposes in poultry production systems and possible dissemination during poultry processing (Van Boeckel et al. 2015) could provide some explanation to the level of resistance detected.

As observed in our study, β-lactamases were detected in 77.6% of 98 isolates screened. This finding explained the molecular basis of the extended-spectrum cephalosporin resistance observed in the *E. coli* isolates recovered from the retail meat products. No detection of β-lactamase genes in 22 isolates, despite phenotypic expression, may suggest mediation by otherβ-lactam resistance mechanisms. Also, considering several of these isolates without β-lactamase genes detected were AmpC phenotypes, chromosomal mutations may also provide some of the explanation to β-lactam resistance observed in this study. In the present study, we detected four different groups of β-lactamases (*bla*_CMY-2_, *bla*_TEM_, *bla*_SHV_, and *bla*_CTXM_ compared to previous studies in Nova Scotia, Canada(Forward et al. 2004), Western Canada (Sheikh et al. 2012), and USA (Doi et al. 2010; Zhao et al. 2012; Sjölund-Karlsson et al. 2013)that reported only *bla*_CMY-2_ and *bla*_TEM_ as the molecular basis for resistance to β-lactams in retail meat products. In addition, the β-lactamase genes detected in the present study have been reported either singly or in combination in chickens(Chalmers et al. 2017), pigs(Kozak et al. 2009; Jahanbakhsh et al. 2016), cattle (Martin et al. 2012; Cormier et al. 2016; Awosile et al. 2018b), dog (Zhang et al. 2018), and as well as in humans in Canada (Denisuik et al. 2013; Awosile et al. 2018a). In all these studies, resistance to ESCs was predominantly mediated by *bla*_CMY-2_, which corroborates the findings of the current study.

While previous studies on retail meat products in Canada did not detect *bla*_CTXM_ genes (Forward et al. 2004; Aslam et al. 2009; Martin et al. 2012; Sheikh et al. 2012; Sjölund-Karlsson et al. 2013), this group of β-lactamases were detected in *E. coli* isolates from retail meat products in the present study. Similarly, in the USA, the *bla*_CTXM_ gene was not reported in retail meat products until 2016 when it was detected in *Salmonella* spp. isolated from retail meats collected as part of National Antimicrobial Monitoring Systems (NARMS)(McDermott et al. 2016). Also, a recent study from NARMS has reported the detection of *bla*_CTXM-14_ and *bla*_CTXM-15_ genes in *E. coli* isolates from retail meat products in the USA(Tadesse et al. 2018). The *bla*_CTXM_ genes are commonly reported from *E. coli* isolated from retail meat products in China (Yu et al. 2015; Li et al. 2016; Wu et al. 2018) and Europe (EFSA Panel on Biological Hazards (BIOHAZ) 2011).The present study and the NARMS studies suggest the possible emergence of *bla*_CTXM_ genes in enteric bacteria isolated from retail meat products in North America. Three different groups of *bla*_CTXM_ genes including *bla*_CTXM-1_, *bla*_CTXM-2_, and *bla*_CTXM-9_ were detected in this study with *bla*_CTXM-1_ and *bla*_CTXM-2_ more commonly detected than *bla*_CTXM-9_. Recent studies in chicken flocks from Ontario (Zhang et al. 2018; Ghosh et al. 2019) and Quebec, Canada (Chalmers et al. 2017) have reported the detection of *bla*_CTXM-1_-like genes similar to this study.

Our results found that poultry-derived *E. coli* isolates were more likely to be carriers of all the three detected groups of β-lactamase genes compared to other retail meat types. This finding is consistent with previous studies (Forward et al. 2004; Sheikh et al. 2012; Zhao et al. 2012; Yu et al. 2015). While 77.6% of the selected ESC-R *E. coli* isolates were positive for at least one β-lactamase gene, some of the *E. coli* isolates (35.7%) were carriers of two or more β-lactamase genes. This carrying capacity for multiple β-lactamase genes in some of the isolates may be responsible for the appearance of cross-resistance to β-lactam antimicrobial class.

ESC-R *E. coli* isolates were recovered in the present study from all the meat product types. A possible explanation for the presence of ESC-resistance in the *E. coli* may be due to the use of ceftiofur in food animal production. A recent report on antimicrobial use in Canada has placed β–lactam antimicrobials, together with tetracyclines, as the most commonly used antimicrobials in food animal production (Ebrahim et al. 2016). In cattle, the use of ceftiofur has been associated with the recovery of ESC-R *E. coli*(Daniels et al. 2009). In Canadian poultry production, use of ceftiofur in broiler flocks has been associated with A2C-resistance (ampicillin, amoxicillin-clavulanate, and cefoxitin) caused by the *bla*_CMY-2_ gene in *E. coli* (Caffrey et al. 2017). While no ESC are registered for use in poultry in Canada, extra-label use in hatcheries is a common practice, especially for the prevention of avian pathogenic *E. coli*(Agunos et al. 2017). In 2013, when the samples for this study were being collected in the Maritimes of Canada, 31% of national poultry flocks were using ceftiofur (Agunos et al. 2017).This extra-label use of ceftiofur may have contributed to the ESC-R observed in this study. However, ceftiofur use has declined steadily over the years following a voluntary withdrawal of use in hatcheries (Dutil et al. 2010; Ebrahim et al. 2016; Agunos et al. 2017). This withdrawal has culminated in a concomitant decline in the prevalence of ESC-R *E. coli* isolates and *Salmonella* Heidelberg recovered from retail chicken, as well as the incidence of infection with ceftiofur-resistant *S*. Heidelberg in humans in Canada (Dutil et al. 2010; Avery et al. 2014).

Environmental contamination of meat carcasses may also provide some explanation for the recovery of ESC-R bacteria. Cross-contamination at the processing plants, poor handling during packaging and transportations of meat carcasses may have contributed to the contamination of retail meat products with ESC-R *E. coli*. Similarly, *bla*_CMY-2_ and *bla*_TEM_ genes detected in *E. coli isolates* from retail meat products in the present study have also been detected in *E. coli* isolates from a commercial beef processing plant in Canada (Aslam et al. 2009). Thus, some of the β-lactamase producing *E. coli* isolated from retail meat may be from contamination of carcasses, their subsequent cuts, and processed meat products present in the processing facility. Contamination of retail meat products with β-lactamase producing bacteria may produce therapeutic consequences in human medicine as these bacteria and/or genes may be transmitted through consumption of improperly cooked meat products. The risk of human exposure to ESC-R *E coli* through consumption of poultry meat products is well established (Depoorter et al. 2012; Evers et al. 2017; Nekouei et al. 2018), therefore food animal production practices that reduce the development and dissemination of antimicrobial resistance, as well as processing plant activities that prevent cross-contamination, are essential strategies to minimize human exposures.

This study has provided information on the prevalence of ESC-R *E. coli* in different retail meat products within the Maritime Provinces, Canada. This study found that using selective culture medium increases the recovery and frequency of ESC-R *E*. coli compared to the traditional culture method. Using molecular techniques, we detected both ESBL and AmpC β-lactamase resistance genes in *E. coli* isolates recovered from retail meat products. We report the detection of *bla*_CTXM_ gene group in *E. coli* isolates from retail meat products for the first time in Canada. ESC-R *E. coli* isolates were recovered more from poultry meat products compared to beef and pork. Similarly, poultry-derived isolates were more likely to be MDR as well as more likely to harbor *bla*_CMY-2_, *bla*_TEM_, and *bla*_CTXM_ resistance genes.

## Acknowledgment

We are grateful to the CIPARS for isolates used in the study were collected as part of routine surveillance within the Maritime Provinces, Canada. Also, we are grateful to Cynthia Mitchell, and Patty McKenna of Atlantic Veterinary College, University of Prince Edward Island Canada for their technical and laboratory support.

